# The genetic and physical interactomes of the *Saccharomyces cerevisiae* Hrq1 helicase

**DOI:** 10.1101/2020.08.28.272674

**Authors:** Cody M. Rogers, Elsbeth Sanders, Phoebe A. Nguyen, Whitney Smith-Kinnaman, Amber L. Mosley, Matthew L. Bochman

## Abstract

The human genome encodes five RecQ helicases (RECQL1, BLM, WRN, RECQL4, and RECQL5) that participate in various processes underpinning genomic stability. Of these enzymes, the disease-associated RECQL4 is comparatively understudied due to a variety of technical challenges. However, *Saccharomyces cerevisiae* encodes a functional homolog of RECQL4 called Hrq1, which is more amenable to experimentation and has recently been shown to be involved in DNA inter-strand crosslink (ICL) repair and telomere maintenance. To expand our understanding of Hrq1 and the RecQ4 subfamily of helicases in general, we took a multi-omics approach to define the Hrq1 interactome in yeast. Using synthetic genetic array analysis, we found that mutations of genes involved in processes such as DNA repair, chromosome segregation, and transcription synthetically interact with deletion of *HRQ1* and the catalytically inactive *hrq1-K318A* allele. Pull-down of tagged Hrq1 and mass spectrometry identification of interacting partners similarly underscored links to these processes and others. Focusing on transcription, we found that *hrq1* mutant cells are sensitive to caffeine and that mutation of *HRQ1* alters the expression levels of hundreds of genes. In the case of *hrq1-K318A*, several of the most highly upregulated genes encode proteins of unknown function whose expression levels are also increased by DNA ICL damage. Together, our results suggest a heretofore unrecognized role for Hrq1 in transcription, as well as novel members of the Hrq1 ICL repair pathway. These data expand our understanding of RecQ4 subfamily helicase biology and help to explain why mutations in human RECQL4 cause diseases of genomic instability.

## INTRODUCTION

A multitude of cellular processes are necessary to ensure the maintenance of genome integrity, including high fidelity DNA replication, recombination and repair, telomere maintenance, and transcription. Among the proteins that are involved, DNA helicases represent one of only a few enzyme classes that are vital to all of these processes (Bochman 2014). Helicases are enzymes that use the power of ATP hydrolysis to drive conformational changes that enable translocation along DNA and unwinding of DNA base pairs (Abdelhaleem 2010; Brosh and Matson 2020). Because these enzymes are involved in so many critical functions *in vivo*, it is unsurprising that mutations in genes encoding helicases are causative of or linked to numerous diseases of genomic instability such as cancer and aging (Monnat 2010; Suhasini and Brosh 2013; Uchiumi *et al*. 2015).

Despite their prominent roles in maintaining genome integrity however, we often lack a detailed understanding of why a particular mutation in a helicase is associated with a pathological disorder. In other words, what cellular processes are impacted that eventually precipitate a disease state when a helicase is mutated? Part of the difficulty in answering this question is that many helicases are multi-functional, and a defect in any one of a number of functions could cause genomic instability (Hickson 2003). Another issue is that helicases are numerous, with > 100 predicted to be encoded by typical eukaryotic genomes (Eki 2010), and many helicases share partially redundant or backup roles, which complicates identification of phenotypes without thorough genomic or proteomic approaches.

One such under-studied and disease-linked helicase is the human RECQL4 protein. Dozens of mutant alleles of *RECQL4* cause three different diseases (Baller-Gerold syndrome (Van Maldergem *et al*. 1993), RAPADILINO (Vargas *et al*. 1992), and Rothmund-Thomson syndrome (Liu 2010)) characterized by a predisposition to cancers, but it is unclear why these mutations cause disease. RECQL4 is difficult to study *in vivo* because it is an evolutionary chimera between a RecQ family helicase and Sld2 (Capp *et al*. 2010), an essential DNA replication initiation factor in lower eukaryotes (Kamimura *et al*. 1998). Helicase activity by RECQL4 is not needed for DNA replication, but pleiotropic defects in replication hamper the analysis of the roles of the helicase domain when studying *recql4* mutants. Similarly, RECQL4 is difficult to study *in vitro* because the protein is large (∼135 kDa) with a natively disordered N-terminus (Keller *et al*. 2014), making the generation of recombinant protein for biochemistry arduous (Macris *et al*. 2006; Bochman *et al*. 2014). Thus, although RECQL4 is reported to be involved in telomere maintenance (Ghosh *et al*. 2011) and DNA inter-strand crosslink (ICL) repair (Jin *et al*. 2008), its mechanism of action in these pathways is unknown.

Recently, we established the *Saccharomyces cerevisiae* Hrq1 helicase as a functional homolog of the helicase portion of RECQL4, showing that it too is linked to telomere maintenance and ICL repair (Bochman *et al*. 2014; Rogers *et al*. 2017; Rogers *et al*. 2020). However, because Sld2 is a separate protein in *S. cerevisiae* and recombinant Hrq1 is more amenable to biochemistry, we have been able to delve into the molecular details of Hrq1 in the maintenance of genome integrity. For instance, Hrq1 synergizes with the helicase Pif1 to regulate telomerase activity, likely establishing telomere length homeostasis *in vivo* (Nickens et al. 2018). In ICL repair, Hrq1 stimulates the translesional nuclease activity of Pso2 to aid in remove of the ICL (Rogers *et al*. 2020). During the course of these investigations, we have also found that alleles of *HRQ1* genetically interact with mutations in the gene encoding the other RecQ family helicase in *S. cerevisiae, SGS1* (Bochman et al. 2014), and that Hrq1 may be involved in the maintenance of DNA motifs capable of forming G-quadruplex (G4) structures (Rogers *et al*. 2017). These facts are mirrored by the interaction of RECQL4 with the human Sgs1 homolog BLM (Singh *et al*. 2012) and the ability of RECQL4 to bind to and unwind G4 DNA (Keller *et al*. 2014).

To gain a more comprehensive understanding of the roles of RecQ4 subfamily helicases in genome integrity, we sought to define the Hrq1 interactome in yeast. Here, we performed synthetic genetic array (SGA) analysis of *hrq1Δ* and *hrq1-K318A* (catalytically inactive mutant) cells using the yeast deletion collection and the temperature-sensitive (TS) collection. Hundreds of significant positive and negative interactions were uncovered, with gene ontology (GO) term enrichment for processes such as transcription and rRNA processing in addition to expected functions such as DNA repair. Mass spectrometry (MS) analysis of proteins that physically interact with Hrq1 returned similar results. Our initial characterization of the link between Hrq1 and transcription revealed that *hrq1* mutant cells are sensitive to the transcription stressor caffeine and that the *hrq1Δ* and *hrq1-K318A* mutations affect the transcription of hundreds of genes, many of which are known or hypothesized to be related to transcription, DNA ICL repair, and the cytoskeleton.

## MATERIALS AND METHODS

### Strain construction

The *HRQ1* gene was deleted in Y8205 (Table 1) by transforming in a NatMX cassette that was PCR-amplified from plasmid pAC372 (a gift from Amy Caudy) using oligonucleotides MB525 and MB526 (Table S1). The deletion was verified by PCR analysis using genomic DNA and oligonucleotides that anneal to regions up- and downstream of the *HRQ1* locus (MB527 and MB528). The confirmed *hrq1Δ* strain was named MBY639. The *hrq1-K318A* allele was introduced into the Y8205 background in a similar manner. First, an *hrq1-K318A(NatMX)* cassette was PCR-amplified from the genomic DNA of strain MBY346 (Bochman *et al*. 2014) using oligonucleotides MB527 and MB528 and transformed into Y8205. Then, genomic DNA was prepared from transformants and used for PCR analyses of the *HRQ1* locus with the same oligonucleotide set to confirm insertion of the NatMX marker. Finally, PCR products of the expected size for *hrq1-K318A(NatMX)* were sequenced using oligonucleotide MB932 to confirm the presence of the K318A mutation. The verified *hrq1-K318A* strain was named MBY644. Hrq1 was tagged with a 3xFLAG epitope in the YPH499 genetic background by transformation of a 3xFLAG(His3MX6) cassette that was PCR-amplified from the pFA6a-3xFLAG-His3MX6 plasmid (Funakoshi and Hochstrasser 2009) using oligonucleotides MB1028 and MB1029. Proper integration was assessed by PCR and sequencing as described above for *hrq1-K318A(NatMX)*. The confirmed Hrq1-3xFLAG strain was named MBY520.

**Table 1.**
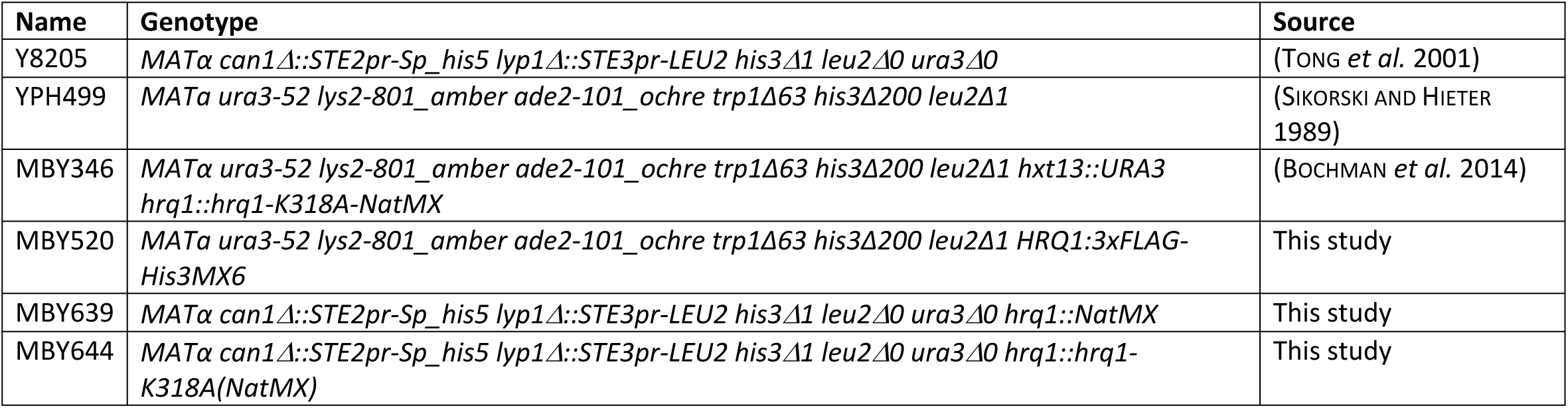
Strains used in this study.

### SGA analysis

SGA analysis of the *hrq1Δ* and *hrq1-K318A* alleles was performed at the University of Toronto using previously described methods (Tong *et al*. 2001; Tong *et al*. 2004). The *hrq1* mutants were crossed to both the *S. cerevisiae* single-gene deletion collection (Giaever and Nislow 2014) and the TS alleles collection (Kofoed *et al*. 2015) to generate double mutants for analysis. Quantitative scoring of the genetic interactions was based on colony size. The SGA score measures the extent to which a double mutant colony size deviates from the colony size expected from combining two mutations together. The data include both negative (putative synthetic sick/lethal) and positive interactions (potential epistatic or suppression interactions) involving *hrq1Δ* and *hrq1-K318A*. The magnitude of the SGA score is indicative of the strength of the interaction. Based on statistical analysis, it was determined that a default cutoff for a significant genetic interaction is *p* < 0.05 and SGA score > |0.08|. It should be noted that only top-scoring interactions were confirmed by remaking and reanalyzing the double mutants by hand.

### Confirmation of top SGA hits

The top five positive and negative interactors with *hrq1Δ* and *hrq1-K318A* from the single-gene deletion and TS arrays were reanalyzed by hand to confirm their phenotypes. Briefly, the SGA query strains MBY639 and MBY644 (Nat^R^) were mated to *MATa* tester strains from the arrays (Kan^R^), sporulated, and then analyzed by random spore analysis (Lichten 2014) and spot dilution growth assays of Nat^R^ Kan^R^ spore clones compared to the parental single-mutant strains and wild-type.

### Hrq1-3xFLAG affinity pulldown

To immunoprecipitate Hrq1-3xFLAG and its associated proteins, strain MBY520 was grown to an optical density at 600 nm (OD_600_) of ∼1.5 in YPD medium at 30°C with shaking. The cells were harvested by centrifugation at 4°C, washed with 50 mL of sterile ice-cold H_2_O, and harvested as before. The cell pellet was then resuspended in 100 μL/g of cells resuspension buffer (20 mM Na-HEPES, pH 7.5, and 1.2% w/v PEG-8000) supplemented with 10 μg/mL DNase I and protease inhibitor cocktail (600 nM leupeptin, 2 μM pepstatin A, 2 mM benzamidine, and 1 mM phenylmethanesulfonyl fluoride). This cell slurry was slowly dripped into liquid nitrogen to generate frozen yeast “popcorn”, which was stored at −80°C until use. To cryo-lyse the cells, the popcorn was ground in a freezer mill with dry ice. The resultant powder was collected into 50-mL conical tubes that were loosely capped and stored at −80°C overnight to allow the dry ice to sublimate away. To perform the Hrq1 pull down, the cell powder was resuspended in 2.5 g powder per 25 mL lysis buffer (40 mM Na-HEPES, pH 7.5, 10% glycerol, 350 mM NaCl, 0.1% Tween-20, and protease inhibitor cocktail) with gentle agitation. Then, 100 U DNase I and 10 μL of 30 mg/mL heparin were added, and the sample was incubated for 10 min at room temperature with gentle agitation. Cellular debris was pelleted by centrifugation at 14,000 x g for 10 min at 4°C. Then, 100 μL of anti-FLAG agarose slurry was washed and equilibrated with lysis buffer, and the clarified lysate and anti-FLAG resin were added to a fresh 50-mL conical tube. This suspension was incubated at 4°C overnight on a nutator. The resin and lysate were subsequently placed in a 30-mL chromatography column, and the lysate was allowed to flow through the resin by gravity. The anti-FLAG agarose was washed with 30 mL lysis buffer, and the beads were then resuspended in 150 μL lysis buffer and transferred to a 1.5-mL microcentrifuge tube. At this point, the sample could be used for proteinase digestion and mass spectrometry analysis, or proteins could be eluted from the resin and examined by SDS-PAGE and Coomassie staining. The untagged control strain (MBY4) was also processed as above to identify proteins that nonspecifically bound to the anti-FLAG agarose.

### Label-free quantitative proteomics interactome analysis

For on-bead digestion, 500 μL of trypsin digestion buffer (50 mM NH_4_HCO_3_, pH 8.5) was used to resuspend the FLAG resin. To this slurry, 10 μL of 0.1 μg/μL Trypsin Gold (Promega) was added and allowed to incubate overnight at 37°C with shaking. After digestion, the FLAG resin was separated from the digested peptides via spin columns and centrifugation. Formic acid (0.1% final concentration) was added to the supernatant to quench the reaction. After digestion, the peptide mix was separated into three equal aliquots. Each replicate was then loaded onto a microcapillary column. Prior to sample loading, the microcapillary column was packed with three phases of chromatography resin: reverse phase resin, strong cation resin, and reverse phase resin, as previous described (Florens and Washburn 2006; Mosley *et al*. 2011; Mosley *et al*. 2013). An LTQ Velos Pro with an in-line Proxeon Easy nLC was utilized for each technical replicate sample, with a 10-step MudPIT method. In MS1, the 10 most intense ions were selected for MS/MS fragmentation, using collision induced dissociation (CID). Dynamic exclusion was set to 90 s with a repeat count of one. Protein database matching of RAW files was performed using SEQUEST and Proteome Discoverer 2.2 (Thermo) against a FASTA database from the yeast Uniprot proteome. Database search parameters were as follows: precursor mass tolerance = 1.4 Da, fragment mass tolerance = 0.8 Da, up to two missed cleavages were allowed, enzyme specificity was set to fully tryptic, and minimum peptide length = 6 amino acids. The false discovery rate (FDR) for all spectra was <1% for reporting as PSM. Percolator, within Proteome Discoverer 2.2, was used to calculate the FDR (Kall *et al*. 2007). SAINT probability scores were calculated as outlined in the Contaminant Repository for Affinity Purification (CRAPome) website (Mellacheruvu *et al*. 2013) and other publications ((Breitkreutz *et al*. 2010; Choi *et al*. 2011; Choi *et al*. 2012; Kwon *et al*. 2013).

### Caffeine sensitivity

The sensitivity of *hrq1* mutant cells to caffeine was assessed both qualitatively and quantitatively. In the first method, cells of the indicated strains were grown overnight in YPD medium at 30°C with aeration, diluted to OD_600_ = 1 in sterile H_2_O, and then serially diluted 10-fold to 10^−4^. Five microliters of these dilutions were then spotted onto YPD agar plates and YPD agar plates supplemented with 10 mM caffeine. The plates were incubated at 30°C for 2 days before capturing images with a flatbed scanner and scoring growth. In the second method, the overnight cultures were diluted to OD_600_ = 0.01 into YPD or YPD supplemented with various concentrations of caffeine. They were then treated as described in (Ononye *et al*. 2020) with slight modifications. Briefly, 200 µL of each culture was placed in duplicate into wells in 96-well plates, and each well was overlaid with mineral oil to prevent evaporation. The plates were incubated (30°C with shaking) in a Synergy H1 microplate reader (BioTek), which recorded OD_660_ measurements at 15-min intervals for 24 h. The mean of the OD_660_ readings for each strain was divided by the mean OD_660_ of the same strain grown in YPD.

### RNA-seq

Cells were harvested from mid-log phase cultures grown in YPD medium, and total RNA was prepared using a YeaStar RNA kit (Zymo Research). Sequencing libraries were prepared, and Illumina sequencing was performed by, Novogene Corporation. Data analysis was then performed by the Indiana University Center for Genomics and Bioinformatics. The sequences were trimmed using the Trim Galore script (https://www.bioinformatics.babraham.ac.uk/projects/trim_galore/), and reads were mapped to the *S. cerevisiae* genome using bowtie2 on local mode (Langmead and Salzberg 2012). Reads were counted, and differential expression analysis were performed using DESeq2 (Love *et al*. 2014). Two or three independent replicates of each strain were analyzed.

### Statistical analysis

Data were analyzed and graphed using GraphPad Prism 6 software. The reported values are averages of ≥ 3 independent experiments, and the error bars are the standard deviation. *P*-values were calculated as described in the figure legends, and we defined statistical significance as *p* < 0.01.

### Data availability

Strains, plasmids, RNA-seq data, and other experimental reagents are available upon request. File S1 contains detailed descriptions of all supplemental files, as well as Table S1 and Figure S1. File S2 contains the full SGA result. File S3 contains the full SAINT analysis results. File S4 contains the transcriptomic changes identified by RNA-seq.

## RESULTS AND DISCUSSION

### The genetic interactome of *HRQ1*

We crossed the *hrq1Δ* and *hrq1-K318A* alleles to the single-gene deletion and TS allele collections to generate all possible double mutants and assessed the growth of the resulting spore clones to identify negative and positive genetic interactions (Tables S2-S5). In total, 117 significant (*p* < 0.05) genetic interactions (76 negative and 41 positive) were identified between *hrq1Δ* and the single-gene deletion collection (Table S2), and 119 (65 negative and 54 positive) were identified between *hrq1Δ* and the TS alleles collection (Table S3). Similarly, 132 significant (*p* < 0.05) genetic interactions (84 negative and 48 positive) were identified between *hrq1-K318A* and the single-gene deletion collection (Table S4), and 102 (41 negative and 61 positive) were identified between *hrq1K318A* and the TS alleles collection (Table S5). When comparing the *hrq1Δ* and *hrq1-K318A* data sets in aggregate, there was ∼39% overlap between the negative genetic interactions (Fig. 1A) and >30% overlap between the positive genetic interactions (Fig. 1B). However, there was very little overlap when comparing negative to positive genetic interactions and *vice versa* (Fig. 1C,D).

**Figure 1.**
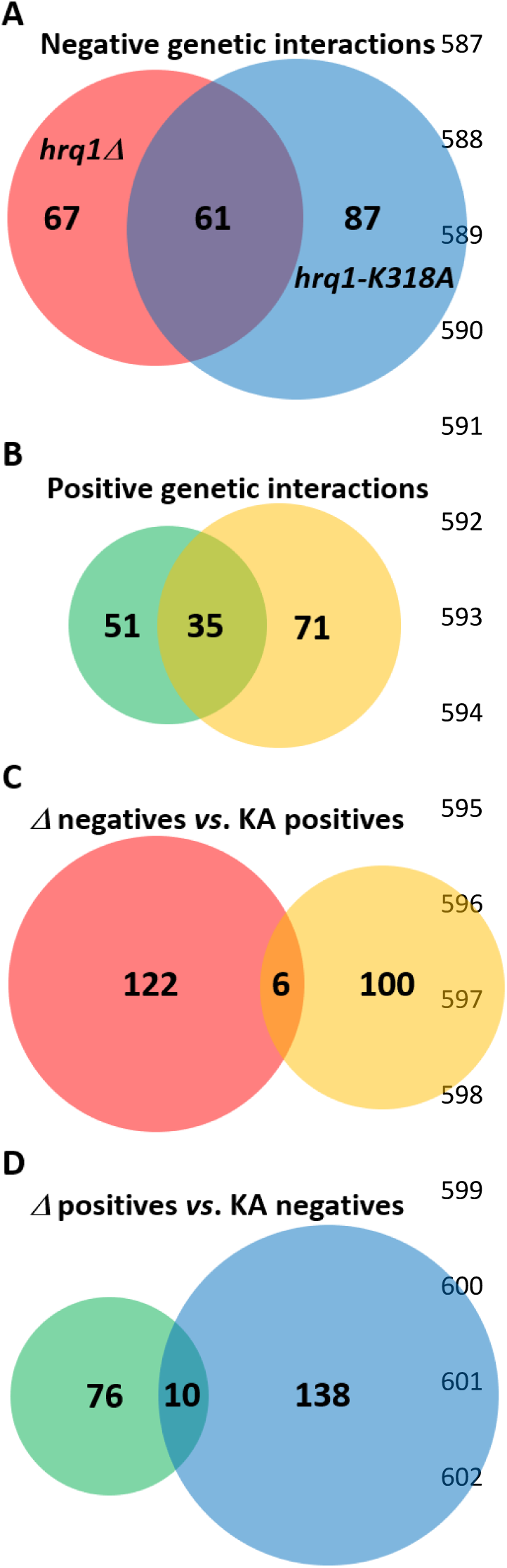
Venn diagrams of the shared synthetic genetic interactions displayed by *hrq1Δ* and *hrq1-K318A*. A) Sixty-one alleles negative interact with both the *hrq1Δ* and *hrq1-K318A* mutations. B) Similarly, 35 alleles positively interact with both the *hrq1Δ* and *hrq1-K318A* mutations. C) Very few of the negative genetic interactors with *hrq1Δ* are common to the set of positive *hrq1-K318A* interactors. D) Likewise, only 10 of the positive genetic interactors with *hrq1Δ* are shared by the set of negative *hrq1-K318A* interactors.

Next, we used GO Term mapping to identify cellular processes enriched for *hrq1* interactors. For all of the negative genetic interactions with *hrq1Δ* and *hrq1-K318A*, the top 10 GO terms were transcription by RNA polymerase II, regulation of organelle organization, DNA repair, chromatin organization, mitotic cell cycle, peptidyl-amino acid modification, cytoskeleton organization, mitochondrion organization, organelle fission, and response to chemical (Table 2). Similarly, for all of the positive genetic interactions with *hrq1Δ* and *hrq1-K318A*, the top 10 GO terms were mitotic cell cycle, cytoskeleton organization, regulation of organelle organization, lipid metabolic process, DNA repair, transcription by RNA polymerase II, chromatin organization, chromosome segregation, organelle fission, and rRNA processing (Table 3).

**Table 2.** Gene Ontology (GO) Term enrichment of negative genetic interactors with *hrq1*.

| GO Term (GO ID) | Genes Annotated to the GO Term | GO Term Usage in Gene List | Genome Frequency of Use |
| --- | --- | --- | --- |
| transcription by RNA polymerase II (GO:0006366) | ABF1, CDC28, CDC73, CEG1, CSE2, EAF7, ESS1, GIM3, HMO1, HTZ1, MED11, NAB3, NUT1, RAD4, SDS3, SGF73, SIN3, SPT15, SPT3, SPT8, SRB2, SRB6, STH1, SUA7, SWI4, TAF11, TAF2, YJR084W | 28 of 191 genes, 14.66% | 536 of 6436 annotated genes, 8.33% |
| regulation of organelle organization (GO:0033043) | APC4, BDF2, CDC15, CDC20, CDC28, CDC73, CTI6, DAM1, EFB1, ESS1, GIC1, LTE1, MOB1, PEF1, PSE1, SDS3, SIN3, SPO16, TGS1, UTH1, VPS41 | 21 of 191 genes, 10.99% | 326 of 6436 annotated genes, 5.07% |
| DNA repair (GO:0006281) | ABF1, ACT1, BDF2, CDC28, CDC73, CST9, EAF7, NSE4, NSE5, POL1, PRP19, RAD14, RAD33, RAD4, RAD52, RAD54, RAD59, RNH201, RTT107, SIN3, STH1 | 21 of 191 genes, 10.99% | 300 of 6436 annotated genes, 4.66% |
| chromatin organization (GO:0006325) | ABF1, BDF2, CDC28, CLP1, CTI6, EAF7, ESS1, GIC1, HTZ1, LGE1, RAD54, SDS3, SGF73, SIN3, SIR1, SPT3, SPT8, STH1, SWC5, UTH1, YCS4 | 21 of 191 genes, 10.99% | 310 of 6436 annotated genes, 4.82% |
| mitotic cell cycle (GO:0000278) | ACT1, APC4, CDC10, CDC15, CDC20, CDC25, CDC28, CDC34, DAM1, GIC1, LTE1, MOB1, PEF1, POL1, PSE1, SIC1, SIN3, SWI4, TUB2, YCS4 | 20 of 191 genes, 10.47% | 373 of 6436 annotated genes, 5.80% |
| peptidyl-amino acid modification (GO:0018193) | ACT1, APJ1, CDC15, CDC28, CDC73, CST9, DBF2, EAF7, ESS1, LIP5, NSE4, NSE5, PSE1, SGF73, SMT3, SPO16, SPT3, SPT8, SWF1, TDA1 | 20 of 191 genes, 10.47% | 244 of 6436 annotated genes, 3.79% |
| cytoskeleton organization (GO:0007010) | ACT1, BBP1, CDC10, CDC15, CDC28, CDC31, CMD1, CTF13, DAM1, EFB1, ENT1, ENT3, GIC1, NDC1, SPC29, STH1, SWF1, TUB2 | 18 of 191 genes, 9.42% | 272 of 6436 annotated genes, 4.23% |
| mitochondrion organization (GO:0007005) | ACT1, ATG1, ATG3, COA4, FCJ1, MDM35, PAM16, PAM17, PHB2, PTC1, QCR2, RCF2, SAM37, TIM18, TOM70, UTH1, YJR120W, YME1 | 18 of 191 genes, 9.42% | 279 of 6436 annotated genes, 4.33% |
| organelle fission<br>(GO:0048285) | APC4, CDC10, CDC15, CDC20, CDC28, CST9, DAM1, DBF2, EBP2, GIC1,<br>LTE1, MOB1, PSE1, RAD52, SPO16, TUB2, YCS4 | 17 of 191<br>genes, 8.90% | 268 of 6436<br>annotated<br>genes, 4.16% |
| response to<br>chemical<br>(GO:0042221) | ACT1, ASK10, GIM3, GPR1, IRA2, MUP3, PTC1, SRB2, TDA1, TIM18,<br>TMA19, TUB2, VPS27, YJR084W, YLR225C, YOS9 | 16 of 191<br>genes, 8.38% | 567 of 6436<br>annotated<br>genes, 8.81% |

**Table 3.** Gene Ontology (GO) Term enrichment of positive genetic interactors with *hrq1*.

| GO Term (GO ID) | Genes Annotated to the GO Term | GO Term Usage in Gene List | Genome Frequency of Use |
| --- | --- | --- | --- |
| mitotic cell cycle<br>(GO:0000278) | ACT1, APC11, BRN1, CDC48, CDC6, CLB3, CSM1, DPB11, IPL1, MCD1, MPS1, MPS3, MYO2, PDS5, PFY1, PSE1, PSF1, SMC4, SPT6, VRP1 | 20 of 151 genes, 13.25% | 373 of 6436 annotated genes, 5.80% |
| cytoskeleton organization<br>(GO:0007010) | ACT1, AIM14, ARP3, CDC48, CLB3, ICE2, IPL1, LAS17, MPS1, MPS2, MPS3, MYO2, NUM1, PFY1, RSP5, SPC29, STH1, TSC11, VRP1 | 19 of 151 genes, 12.58% | 272 of 6436 annotated genes, 4.23% |
| regulation of organelle organization<br>(GO:0033043) | AIM14, APC11, ARP3, CDC48, CDC6, CLB3, IPL1, LAS17, MPS1, PCP1, PFY1, PSE1, RSP5, SEC23, SGV1, SPT6, TSC11, VRP1 | 18 of 151 genes, 11.92% | 326 of 6436 annotated genes, 5.07% |
| lipid metabolic process<br>(GO:0006629) | ALG14, CDC1, CHO2, DGA1, GAA1, GPI10, GPI12, GPI2, GWT1, LCB1, MGA2, OPI3, PHS1, RSP5, SAC1, SUR1, TSC11, VPS4 | 18 of 151 genes, 11.92% | 347 of 6436 annotated genes, 5.39% |
| DNA repair<br>(GO:0006281) | ACT1, ARP8, CDC1, DPB11, IXR1, MCD1, NHP10, PDS5, POB3, POL3, PSF1, RAD3, RNH201, RSC2, SLX5, SLX8, STH1, TEL1 | 18 of 151 genes, 11.92% | 300 of 6436 annotated genes, 4.66% |
| transcription by RNA polymerase II<br>(GO:0006366) | CAM1, CSE2, IXR1, MGA2, MOT1, NHP10, PDC2, POB3, RAD3, RGR1, RSC2, RSP5, SGV1, SPT6, STH1 | 15 of 151 genes, 9.93% | 536 of 6436 annotated genes, 8.33% |
| chromatin organization<br>(GO:0006325) | ARP8, CAC2, CDC6, IES1, MGA2, MPS3, NHP10, ORC6, POB3, RSC2, RSP5, SPT6, STH1, TEL1 | 14 of 151 genes, 9.27% | 310 of 6436 annotated genes, 4.82% |
| chromosome segregation<br>(GO:0007059) | APC11, BRN1, CDC48, CSM1, IPL1, MCD1, MPS1, MPS3, PDS5, RSC2, SMC4, SPC24, STH1 | 13 of 151 genes, 8.61% | 210 of 6436 annotated genes, 3.26% |
| organelle fission<br>(GO:0048285) | APC11, BRN1, CLB3, CSM1, IPL1, MCD1, MPS1, MPS3, NUM1, PDS5, PSE1, SMC4 | 12 of 151 genes, 7.95% | 268 of 6436 annotated genes, 4.16% |
| rRNA processing<br>(GO:0006364) | BMS1, FAL1, MAK5, MOT1, MRM2, POP4, RPF2, RPS23A, RPS6B, RPS9B, RSP5, SLX9 | 12 of 151 genes, 7.95% | 352 of 6436 annotated genes, 5.47% |

A discussion of the strongest negative synthetic genetic interactions with *hrq1Δ* and *hrq1-K318A* is included in Sanders *et al*. (companion paper)^2^ Briefly, this included synthetic interactions with genes encoding genome integrity factors (*e*.*g*., *RAD14* and *CBC2*) and mitochondrial proteins (*e*.*g*., *MRM2* and *TOM70*), consistent with the known roles of Hrq1 and human RECQL4 in genome maintenance (Ghosh *et al*. 2011; Singh *et al*. 2012; Choi *et al*. 2013; Bochman *et al*. 2014; Choi *et al*. 2014; Leung *et al*. 2014; Rogers *et al*. 2017; Nickens *et al*. 2018; Rogers *et al*. 2020) and their nuclear and mitochondrial localization (Croteau *et al*. 2012; Koh *et al*. 2015; Kumari *et al*. 2016).

Deletion alleles of *ARP8* and *SHE1* and TS alleles of *ACT1, ARP3, CSE2, MPS1*, and *MPS3* are among the strongest positive synthetic genetic interactors with *hrq1Δ* and/or *hrq1-K318A* (Tables S3 and S5). Arp8 is a chromatin remodeling factor (Shen *et al*. 2000), and Cse2 is a Mediator complex subunit required for RNA polymerase II regulation (Gustafsson *et al*. 1998), consistent with the GO Term enrichment described above. This may suggest that like human RECQL5 (Aygun *et al*. 2008; Izumikawa *et al*. 2008; Saponaro *et al*. 2014), Hrq1 plays a role in transcription.

She1 is a microtubule-associated protein (Bergman *et al*. 2012), as is human RECQL4 (Yokoyama *et al*. 2019). Likewise, Mps1 and Mps3 are linked to the microtubule cytoskeleton as proteins necessary for spindle pole body function (Friederichs *et al*. 2011; Meyer *et al*. 2013). We attempted to determine if Hrq1 also binds to microtubules using an *in vitro* microtubule co-sedimentation assay (Walker *et al*. 2019), but found that Hrq1 alone pellets during ultracentrifugation (data not shown). We hypothesize that this is due to the natively disordered N-terminus of Hrq1 (Rogers *et al*. 2017; Rogers *et al*. 2020), which may mediate liquid-liquid phase separation (LLPS) of recombinant Hrq1 in solution. Ongoing experiments are addressing the LLPS of Hrq1 alone and in combination with its ICL repair cofactor Pso2 (Rogers *et al*. 2020).

*ACT1* encodes the *S. cerevisiae* actin protein (Gallwitz and Seidel 1980), and Arp3 is a subunit of the Arp2/3 complex that acts as an actin nucleation center (Machesky and Gould 1999). It is unclear why mutation of these cytoskeletal factors yields increased growth in combination with *hrq1* mutations. However, *arp3* mutation also decreases telomere length (Ungar *et al*. 2009). Thus, this synthetic genetic effect may be related to the role of Hrq1 in telomere maintenance (Bochman *et al*. 2014; Nickens *et al*. 2018).

### The physical interactome of Hrq1

To complement our genetic analysis of *hrq1* alleles, we also sought to identify the proteins that physically interact with Hrq1 *in vivo*. To do this, we cloned the sequence for a 3xFLAG tag in frame to the 3’ end of the *HRQ1* gene, replacing its native stop codon. The tag does not disrupt any known activities of Hrq1, as demonstrated by the DNA ICL resistance of the Hrq1-3xFLAG strain (Fig. S1 and data not shown). Next, we snap-froze and cryo-lysed cells to preserve macromolecular complexes in near-native states (Mosley *et al*. 2011), immunoprecipitated Hrq1-3xFLAG and its associated proteins from the lysates, and analyzed them using a quantitative proteomics approach.

Overall, 290 interacting proteins were identified (Table S6), 77 of which had a SAINT score > 0.75 and were thus considered significant (Fig. 2A). These 77 proteins are enriched for GO Term processes such as rRNA processing, ribosomal small subunit biogenesis, ribosomal large subunit biogenesis, cytoplasmic translation, transcription by RNA polymerase I, transcription by RNA polymerase II, RNA modification, DNA repair, chromatin organization, and peptidyl-amino acid modification (Table 4). Further, these categories are representative of the entire set of 290 proteins.

**Table 4.** Gene Ontology (GO) Term enrichment of proteins that physically interact with Hrq1.

| GO Term (GO ID) | Genes Annotated to the GO Term | GO Term Usage in Gene List | Genome Frequency of Use |
| --- | --- | --- | --- |
| rRNA processing<br>(GO:0006364) | BMS1, BUD21, CBF5, CIC1, DBP10, DBP3, DBP9, DHR2, DIP2, ECM16, ENP2, ERB1, ESF1, FUN12, GAR1, KRE33, MAK5, MOT1, MPP10, MRD1, NHP2, NOP56, NOP8, NSA2, NSR1, NUG1, RLP7, ROK1, RPL1A, RPS6A, RPS8A, RRP12, RRP8, TSR1, URB1, UTP10, UTP22, UTP9 | 38 of 75 genes,<br>50.67% | 352 of 6436<br>annotated<br>genes, 5.47% |
| ribosomal small subunit biogenesis<br>(GO:0042274) | BMS1, BUD21, DHR2, DIP2, ECM16, ENP2, FUN12, KRE33, MPP10, MRD1, NSR1, ROK1, RPS19A, RPS6A, RPS8A, RRP12, SGD1, TSR1, UTP10, UTP22, UTP9 | 21 of 75 genes,<br>28.00% | 149 of 6436<br>annotated<br>genes, 2.32% |
| ribosomal large subunit biogenesis<br>(GO:0042273) | CIC1, DBP10, DBP3, DBP9, ERB1, MAK5, NHP2, NOC2, NOP8, NSA2, NUG1, RIX7, RLP7, RPL12A, RPL1A, RRP8, SDA1, URB1 | 18 of 75 genes,<br>24.00% | 122 of 6436<br>annotated<br>genes, 1.90% |
| cytoplasmic translation<br>(GO:0002181) | FUN12, NIP1, RPL12A, RPL1A, RPL23A, RPL43A, RPS19A, RPS25B, RPS6A, RPS8A | 10 of 75 genes,<br>13.33% | 201 of 6436<br>annotated<br>genes, 3.12% |
| transcription by RNA polymerase I<br>(GO:0006360) | CDC73, CTR9, DHR2, LEO1, MOT1, RPA49, RPB5, UTP10, UTP9 | 9 of 75 genes,<br>12.00% | 69 of 6436<br>annotated<br>genes, 1.07% |
| transcription by RNA polymerase II<br>(GO:0006366) | CDC73, CTR9, HHF1, HTA1, LEO1, MOT1, RPB5, RTG3 | 8 of 75 genes,<br>10.67% | 536 of 6436<br>annotated<br>genes, 8.33% |
| RNA modification<br>(GO:0009451) | AIR2, CBF5, GAR1, KRE33, NHP2, NOP56, PUS1, RRP8 | 8 of 75 genes,<br>10.67% | 177 of 6436<br>annotated<br>genes, 2.75% |
| DNA repair<br>(GO:0006281) | CDC73, CTR9, HTA1, LEO1, PDS5, RFC3 | 6 of 75 genes,<br>8.00% | 300 of 6436<br>annotated<br>genes, 4.66% |
| chromatin organization<br>(GO:0006325) | CTR9, FPR3, FPR4, HHF1, HTA1, LEO1 | 6 of 75 genes,<br>8.00% | 310 of 6436<br>annotated<br>genes, 4.82% |
| peptidyl-amino acid modification<br>(GO:0018193) | CDC73, CTR9, FPR3, FPR4, HHF1, LEO1 | 6 of 75 genes,<br>8.00% | 244 of 6436<br>annotated<br>genes, 3.79% |

**Figure 2.**
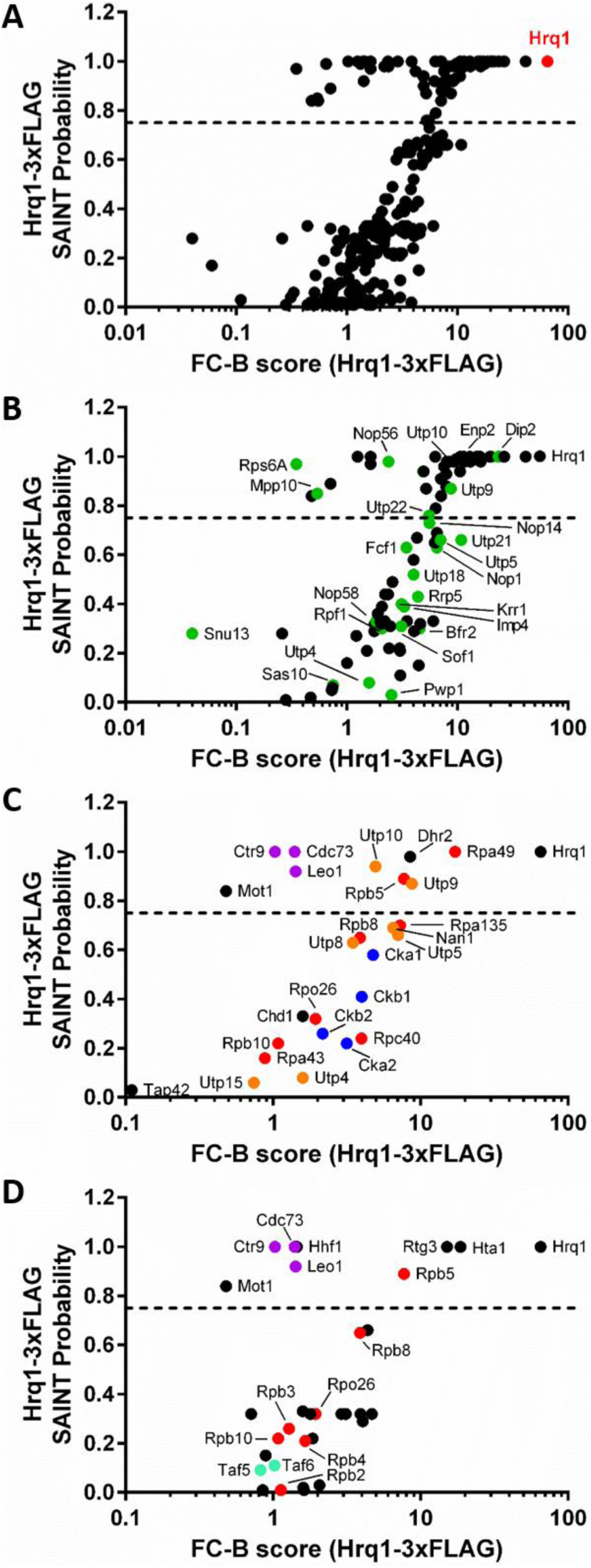
Identification of the Hrq1-3xFLAg interactome by IP-MS and SAINT. A) Overview of the 290 interactions identified by SAINT in anti-FLAG Hrq1 purifications. The graph compares the FC-B score against the SAINT probability score. The dashed line represents the 0.75 probability cut-off. The highest confidence hit, Hrq1, is shown in red. Subsets of the 290 interactors enriched for rRNA processing and ribosomal small subunit biogenesis (B), transcription by RNA polymerase I (C), and transcription by RNA polymerase II (D) factors are also shown. Members of macromolecular complexes associated with these processes are labeled and color coded: small ribosomal subunit processome (https://www.yeastgenome.org/complex/CPX-1604), green; RNA polymerase I (https://www.yeastgenome.org/complex/CPX-1664), II (https://www.yeastgenome.org/complex/CPX-2662), and III (https://www.yeastgenome.org/go/GO:0005666), red; PAF1 complex (https://www.yeastgenome.org/complex/CPX-1726), purple; casein kinase 2 (https://www.yeastgenome.org/complex/CPX-581), blue; UTP-A complex (https://www.yeastgenome.org/complex/CPX-1409), orange; and TFIID (https://www.yeastgenome.org/complex/CPX-1642), teal. All identifiers for these data are included in Table S6.

To demonstrate the robustness of these data, we identified Hrq1-interacting proteins that are subunits of larger macromolecular complexes involved in several of the GO Term processes listed above. For instance, among the rRNA processing and ribosomal small subunit biogenesis proteins (Fig. 2B), several members of the small ribosomal subunit processome (https://www.yeastgenome.org/complex/CPX-1604) are significant Hrq1 interactors. Many more such proteins had SAINT scores < 0.75, suggesting that they may be secondary interactors (*i*.*e*., they physically interact with a significant Hrq1 interactor rather than Hrq1 directly) and/or more weakly associated subunits of the processome. Similarly, the transcription by RNA polymerase I (Fig. 2C) and transcription by RNA polymerase II (Fig. 2D) proteins contain members of multiple macromolecular complexes, including the RNA polymerase I (https://www.yeastgenome.org/complex/CPX-1664) and RNA polymerase II (https://www.yeastgenome.org/complex/CPX-2662) complexes themselves. As with the SGA data above, these links to transcription are intriguing and reminiscent of the links of human RECQL5 to transcription (Aygun *et al*. 2008; Izumikawa *et al*. 2008; Saponaro *et al*. 2014).

### Transcriptomic perturbations caused by mutation of *HRQ1*

Due to the links between the Hrq1 and transcription identified through SGA and immunoprecipitation-mass spectrometry (IP-MS), we decided to determine if the *S. cerevisiae* transcriptome is altered by *HRQ1* mutation. First, we tested the effects of the general transcription stressor caffeine (Kuranda *et al*. 2006) on *hrq1Δ* and *hrq1-K318A* cells. As shown in Figure 3A, the *hrq1-K318A* strain was much more sensitive to 10 mM caffeine than wild-type, though the *hrq1Δ* strain displayed little-to-no caffeine sensitivity. To obtain more quantitative data, we performed growth curve experiments for wild-type, *hrq1Δ*, and *hrq1-K318A* cells in the absence and presence of increasing concentrations of caffeine. At high levels of caffeine, the *hrq1Δ* strain was significantly (*p* < 0.0001) more sensitive than wild-type, but again, the *hrq1-K318A* mutant displayed greater sensitivity at a wider range of concentrations (Fig. 2B). These data mirror the increased sensitivity of the *hrq1-K318A* strain to DNA ICL damage compared to the *hrq1Δ* mutant (Bochman *et al*. 2014; Rogers *et al*. 2020), suggesting that the Hrq1-K318A protein is still recruited to its sites of action *in vivo* but somehow disrupts transcription as a catalytically inert roadblock.

**Figure 3.**
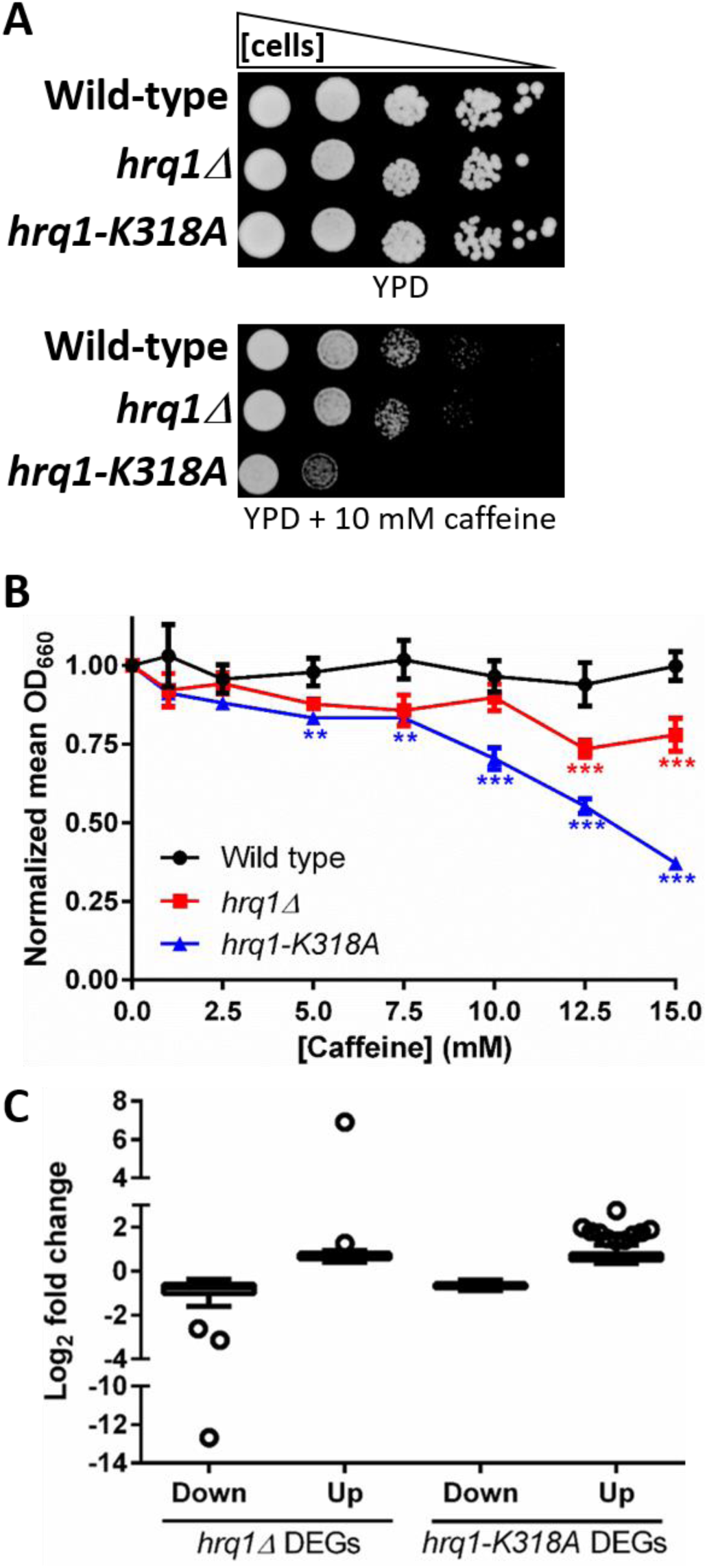
Mutation of *HRQ1* affects transcription. A) Tenfold serial dilutions of the indicated strains on rich medium (YPD) and YPD containing 10 mM caffeine. The *hrq1-K318A* cells are more sensitive to caffeine than the mild sensitivity displayed by the *hrq1Δ* mutant. B) Quantitative analysis of the effects of caffeine on the growth of *hrq1* cells. The normalized values were averaged from ≥ 3 independent experiments and compared to wild-type growth at the same caffeine concentration by one-way ANOVA. **, *p* < 0.001 and ***, *p <* 0.0001. As in (A), *hrq1Δ* cells display milder sensitivity to caffeine than *hrq1-k318A* cells. C) Analysis of the distribution of the magnitudes of expression changes of the DEGs. The log_2_ fold change data for the significantly downregulated (Down) and upregulated (Up) DEGs in *hrq1Δ* and *hrq1-K318A* cells compared to wild-type cells are shown as box and whisker plots drawn using the Tukey method. The individually plotted points outside of the inner fences represent outliers (*i*.*e*., expression changes with the largest absolute values) and correspond to genes whose log_2_ fold change value is less than the value of the 25^th^ quartile minus 1.5 times the inter-quartile distance (IQR) for downregulation or genes whose log_2_ fold change value is greater than the value of the 75^th^ quartile plus 1.5IQR for upregulation.

To gain a transcriptome-wide perspective, we also performed RNA-seq analysis of wild-type, *hrq1Δ*, and *hrq1-K318A* cells. Compared to wild-type, 107 genes were significantly downregulated and 28 genes were significantly upregulated in *hrq1Δ* cells (Table S7). Similarly, 301 and 124 genes were down- and upregulated, respectively, in *hrq1-K318A* cells compared to wild-type. Similar to the SGA and proteomic data sets, the GO Terms of these differentially expressed genes (DEGs) were enriched for processes such as response to chemical, meiotic cell cycle, mitotic cell cycle, rRNA processing, and chromosome segregation (Table S8).

Figure 3C shows the frequency distribution of all of the changes in expression in the *hrq1* cells compared to wild-type, separated by down- and upregulated DEGs for each mutant. Outliers are denoted as single points, representing the transcripts whose abundances changed the most. The expression changes in most DEGs were mild decreases or increases, but several varied greatly from wild-type. As an internal control, we found that the transcription of *HRQ1* in *hrq1Δ* cells displayed the largest decrease among all data sets relative to wild-type (Fig. 3C).

The largest number of outliers were the 10 upregulated DEGs in *hrq1-K318A* cells. These included genes encoding two cell wall mannoproteins (*TIP1* and *CWP1*) (van der Vaart *et al*. 1995; Fujii *et al*. 1999), a heat shock protein (*HSP30*) (Piper *et al*. 1997), a protein required for viability in cells lacking mitochondrial DNA (*ICY1*) (Dunn and Jensen 2003), a predicted transcription factor whose nuclear localization increases upon DNA replication stress (*STP4*) (Tkach *et al*. 2012), a protein of unknown function whose levels increase in response to replication stress (*YER053C-A*), a factor whose over-expression blocks cells in G1 phase (*CIP1*) (Ren *et al*. 2016), and three proteins of unknown function that are induced by ICL damage (*YLR297W, TDA6*, and *FMP48*) (Dardalhon *et al*. 2007). The latter are particularly tantalizing considering the known function of Hrq1 in ICL repair (Bochman 2014; Rogers *et al*. 2017; Rogers *et al*. 2020). Perhaps the *YLR297W, TDA6*, and *FMP48* gene products function in the Hrq1-Pso2 ICL repair pathway, and their levels must be elevated to compensate for the catalytically crippled Hrq1-K318A mutant. Alternatively, they may represent members of a back-up ICL repair pathway that is activated when the Hrq1-Pso2 pathway is ablated. In either case, it should be noted that the RNA-seq experiments were performed in the absence of exogenous ICL damage, but the *hrq1-K318A* cells appear already primed to deal with ICLs in the absence of functional Hrq1. The reasons for this are currently unknown, but our ongoing experiments are addressing this phenomenon.

### Conclusions and perspectives

Here, we used a multi-omics approach to comprehensively determine the *S. cerevisiae* Hrq1 interactome. The data reported here and in our companion manuscript (Sanders *et al*., companion paper) greatly expand the known genetic and physical interaction landscape of Hrq1 in yeast, including synthetic genetic interactions with and transcriptomic changes caused by the strong *hrq1-K318A* allele. Various links to the known and putative roles of Hrq1 and its homologs in DNA repair, telomere maintenance, and the mitochondria were found, as well as novel connections to the cytoskeleton and transcription.

Our concurrent data also indicate that the second *S. cerevisiae* RecQ family helicase, Sgs1, is also involved in transcription (Sanders *et al*., companion paper). However, it is unclear if Hrq1 and Sgs1 act together during transcription or have distinct roles, and it is unknown what these roles are. Human RECQL5 physically interacts with RNA polymerase II, controlling transcription elongation (Saponaro *et al*. 2014). It may also function at the interface of DNA repair and transcription by helping to resolve replication-transcription conflicts (Hamadeh and Lansdorp 2020). It is reasonable to hypothesize that Hrq1 and/or Sgs1 function similarly and, in the case of Hrq1, perhaps in the transcription-coupled repair of DNA ICL lesions. Future work should address these hypotheses, as well as the others raised throughout this manuscript, to further characterize the roles of RecQ helicases in the maintenance of genome integrity. Similar to the mechanistic identification of the roles of Hrq1 in yeast (Bochman 2014; Nickens *et al*. 2018; Rogers *et al*. 2020), we anticipate that these data will spur additional research into exciting and unexpected functions of RecQ4 subfamily helicases.

## ACKNOWLEDGEMENTS

We thank Amy Caudy for sharing plasmids, the University of Toronto for performing the SGA analyses, Michael Costanzo and members of the Boone lab for help with data collection and interpretation, and members of the Bochman and Mosley labs for critically reading this manuscript. This research was supported by the College of Arts and Sciences, Indiana University (to MLB), the Indiana University Collaborative Research Grant fund of the Office of the Vice President for Research (to MLB and ALM), the American Cancer Society (RSG-16-180-01-DMC to MLB), and the National Institutes of Health (1R35GM133437 to MLB).

## FOOTNOTES

^1^ Sanders *et al*., Comprehensive synthetic genetic array analysis of alleles that interact with mutation of the *Saccharomyces cerevisiae* RecQ helicases Hrq1 and Sgs1, <u>submitted as a companion paper to *G3*</u>.

## SUPPLEMENTAL MATERIALS

**Table S1.**
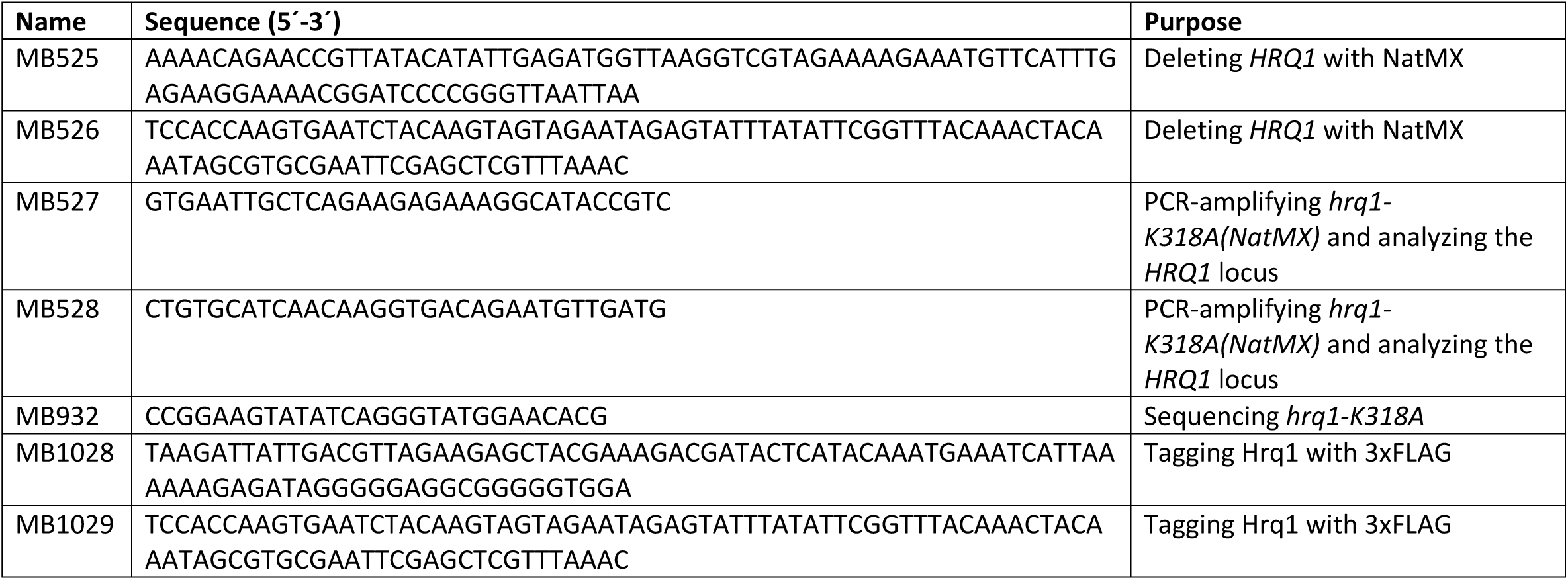
Oligonucleotides used in this study.

**Table S2**. Negative genetic interactions with *hrq1Δ* (see file S2).

**Table S3**. Positive genetic interactions with *hrq1Δ* (see file S2).

**Table S4**. Negative genetic interactions with *hrq1-K318A* (see file S2).

**Table S5**. Positive genetic interactions with *hrq1-K318A* (see file S2).

**Table S6**. Hrq1-interacting proteins (see file S3).

**Table S7**. Gene expression levels in *hrq1Δ* and *hrq1-K318A* cells compared to wild-type (see file S4).

**Figure S1.**
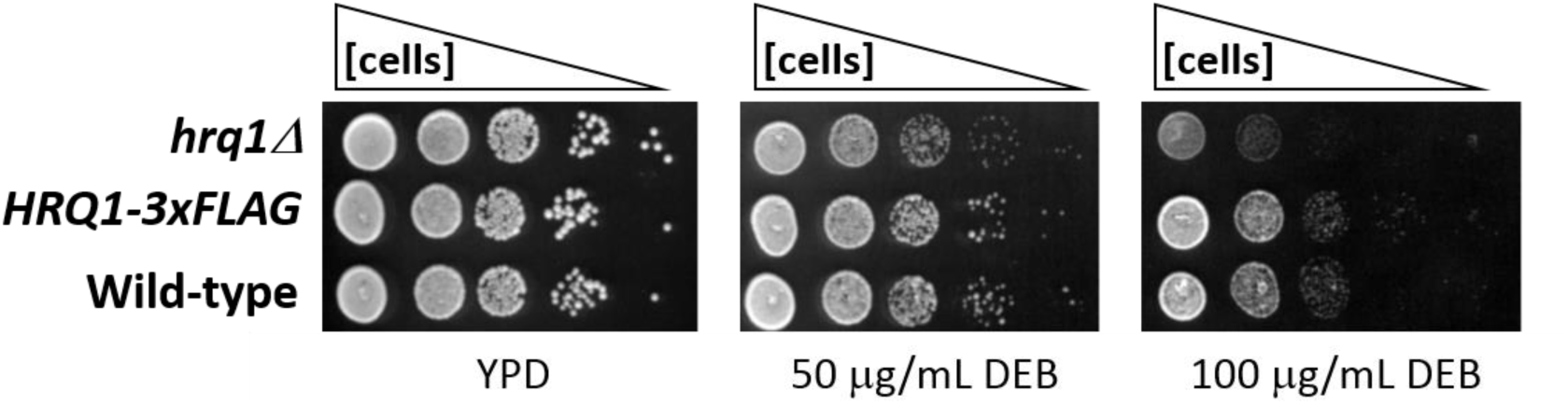
The C-terminal 3xFLAG tag does not interfere with the role of Hrq1 in DNA inter-strand crosslink repair. The indicated strains were grown overnight in YPD medium at 30°C with aeration, diluted to OD_600_ = 1 in sterile H_2_O, and then serially diluted 10-fold to 10^−4^. Five microliters of these dilutions were then spotted onto YPD agar plates and YPD agar plates supplemented with 50 or 100 μg/mL diepoxybutane (DEB). The plates were incubated at 30°C for 2 days before capturing images with a flatbed scanner and scoring growth.

## Footnotes

2 Sanders *et al*., Comprehensive synthetic genetic array analysis of alleles that interact with mutation of the *Saccharomyces cerevisiae* RecQ helicases Hrq1 and Sgs1, <u>submitted as a companion paper to *G3*</u>.

